# Single-molecule validation and optimized protocols for the use of secondary nanobodies in multiplexed immunoassays

**DOI:** 10.1101/2025.02.28.640765

**Authors:** Rebecca S. Saleeb, Judi O’Shaughnessy, Ryan Ferguson, Candace T. Adams, Mathew H. Horrocks

## Abstract

Recently developed secondary nanobodies or single-domain antibodies present a powerful tool for immunodetection. Unlike traditional antibodies, their monovalence enables pre-association with primary antibodies prior to sample staining, presenting a straightforward affinity-based antibody labeling solution. This not only simplifies and streamlines immunoassays, it also supports multiplexed techniques where conflicts in the species of the desired primary antibodies preclude standard indirect immunostaining. Despite these advantages, the use of secondary nanobodies remains sparse, due perhaps to a lack of evaluation on their suitability for assays requiring quantification and an assessment of optimal protocols for their use. Here, we present a set of experiments spanning single-molecule detection to cell imaging that can be used to validate secondary nanobody binding, specificity, and their propensity for mis-targeted binding in multiplex assays. Using these tools, we analyzed the binding properties of commercially available secondary nanobodies and outline optimized protocols for their robust use.

---

A long-standing challenge for immunofluorescence (IF) assays is the detection of multiple target molecules while maintaining target specificity. The number of targets that can be probed is traditionally constrained by the number of different fluorophores that can be spectrally separated. However, approaches to overcome this through cyclical imaging or spectral unmixing have been long established^1,2^. Recent advances in automation^3^ and label exchange methodology^4–10^ now enable more than 50 markers to be detected *in situ*^11^. This has aided progress towards imaging-based spatial proteomics, but uptake and implementation has been slow, likely in-part due to increased sample preparation complexities.

A particular bottleneck is the generation of uniquely labeled primary antibodies (IgG). Dependent on the strategy used, this process can be costly, resource greedy, can have low efficiency, suffer batch variability, may impact binding kinetics, and commercially supplied antibodies may be unsuitable dependent on buffer composition. Indirect immunostaining can instead de-couple the primary antibody, which targets the protein of interest, from the detection molecule via a secondary antibody. This enables a wide range of detection strategies, such as fluorophores, chromogens, biotin-mediated and DNA-based labels, to be utilized without reliance on direct conjugation to the primary antibody. The notable limitation of indirect immunostaining is that multiplexed experiments must employ primary antibodies of different species or classes to enable species-specific targeting of the secondary antibody. Pre-incubating the primary and secondary to couple them *prior* to sample staining could overcome this, however full-size secondary antibodies drive antibody clustering due to their bivalence.

Secondary nanobodies (2.Nbs) are an intriguing solution to this. Nanobodies (Nbs), also known as VHH or single-domain antibodies (sdAbs), are the isolated variable domain of heavy chain-only antibodies (HCAbs) uniquely found in camelids^12^ and cartilaginous fish^13^. They have advantageous properties over conventional IgG for immuno-based techniques due to their 10-fold smaller size, greater stability, reduced immunogenicity, strong antigen binding affinity, lower production cost, and straight forward engineering^14,15^. Of particular interest for multiplex imaging is their monovalence, which enables pre-mixing with the primary antibody for single-step indirect immunostaining without generating antibody clusters^14,15^. This powerful approach allows even small quantities of primary antibodies to be quickly and easily pre-labeled with any probe prior to staining for total experimental flexibility without species complexity. This has shown recent promise for the simplification of highly multiplexed imaging workflows^16–18^ and could offer a low-cost solution to standard 2-3 channel IF assays where preferred antibodies present a species clash.

Despite this opportunity, uptake of 2.Nbs for indirect immunostaining has been slow. We speculate this stems from concerns that antibody “hopping” and incomplete labeling may produce misleading results (schematized in Figure 1A). Quantitative analysis of this effect is critical to the integrity of future studies employing these tools. Though encouraging multiplexed cell images have been qualitatively presented^14,15^, no quantitative analysis has been carried out to assess this concern. To address this, we present here simple methodologies and analysis pipelines for the assessment of 2.Nb cross-over staining. Through the implementation of these assays, we show that pre-labeling can be done robustly, and we provide optimized and validated protocols for the use of 2.Nbs in the detection of targets from the cell to the single molecule level.

**Figure 1.**
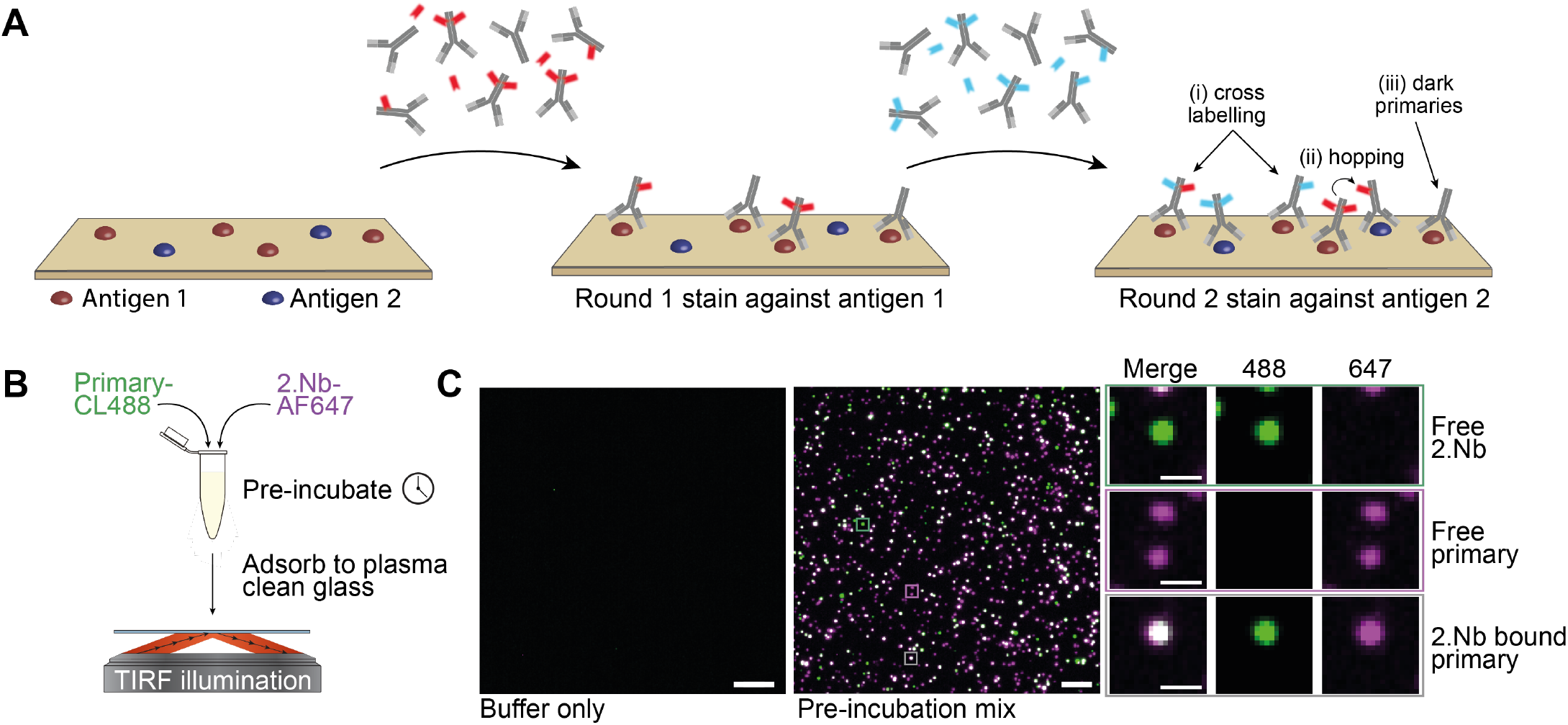
A single-molecule assay to validate secondary Nb binding efficiency. (**A**) Schematic representation of potential artefacts generated by one-step 2.Nb-based indirect immunofluorescence with primary IgGs raised in the same species. Antibodies (gray) and nanobodies against antigen 1 (red) or antigen 2 (blue) are pre-mixed prior to staining. Incomplete primary IgG/2.Nb binding may produce invisible or dark primaries and mis-targeted cross-labeling. (**B**) Schematic representation of binding validation assay that adsorbs pre-mixed fluorescently labeled primary IgG (green) and 2.Nb (magenta) on to plasma treated coverglass for TIRFM analysis. (**C**) Example TIRFM images captured using the assay in (B) when buffer only (left) or pre-mixed antibodies present (centre), scale bars 5 µm. Free 2.Nb (green), free primary IgG (magenta), and bound primary IgG/2.Nb (white) are visible. Single channel images shown (right), scale bars 1 µm.

## Results

### Single-molecule characterization of 2.Nb binding efficiency

The interaction between an antibody and its target antigen is non-covalent and thus reversible. Greater affinity and stability is characterized by a high association rate (k_on_) and low dissociation rate (k_off_) and thus varies between immunoglobulin clones. A sub-optimal k_on_ will result in lower target engagement, whereas sub-optimal k_off_ will result in a short duration of engagement. In these cases, secondary detection antibodies may fail to occupy all binding sites, producing unlabeled or ‘dark’ primaries (Figure 1A (iii)). This can reduce detection sensitivity, but is not prohibitive for traditional indirect immunofluorescence. However, when using pre-mixed primary IgG and 2.Nbs for multiplexing primary antibodies of the same species, incomplete labeling leaves both primary IgG epitope sites and free 2.Nb available for off-target ‘cross-labeling’ (Figure 1A (i)). In addition, if the k_off_ of bound 2.Nbs is not negligible across the staining and detection timeline, Nb ‘hopping’ may occur where the secondary Nb moves from an on-target association to an off-target association (Figure 1A (ii)).

To assess the presence of these artefacts and their contribution during one-step IF, we designed a single-molecule assay in which primary IgG that is directly conjugated to CoraLite Plus 488 (CL488) is pre-mixed with an Alexa Fluor 647 (AF647)-conjugated 2.Nb, and subsequently adsorbed to argon plasma-treated cover glass at single-molecule density (Figure 1B). The composition of the mixture was analyzed by total internal fluorescence microscopy (TIRFM), where AF647-only puncta indicate free 2.Nb, CL488-only puncta indicate free primary IgG, and coincident CL488 and AF647 puncta indicate 2.Nb-bound primary IgG (Figure 1C).

Using this binding assay, we analyzed the degree of 2.Nb target engagement as its concentration relative to the primary IgG increased. Due to the symmetrical nature of primary IgG, each antibody has two nanobody binding sites. It is therefore expected that a minimum two-fold excess of monoclonal 2.Nb is required to saturate all sites. Further excess, however, may help shift the equilibrium towards the bound state. Published work and vendor protocols have proposed variable primary IgG to 2.Nb ratios to date, ranging from 1.3-3.8 molar excess of the 2.Nb^16,18^. However, in the case of the 10E10 anti-rabbit IgG 2.Nb clone, we observed that the percentage of primary IgG associated with the nanobody reached a plateau of 82 ± 2.82% and further 2.Nb excess only boosted the free Nb population (Figure 2A-B), increasing the potential for non-specific associations in multiplexed assays.

**Figure 2.**
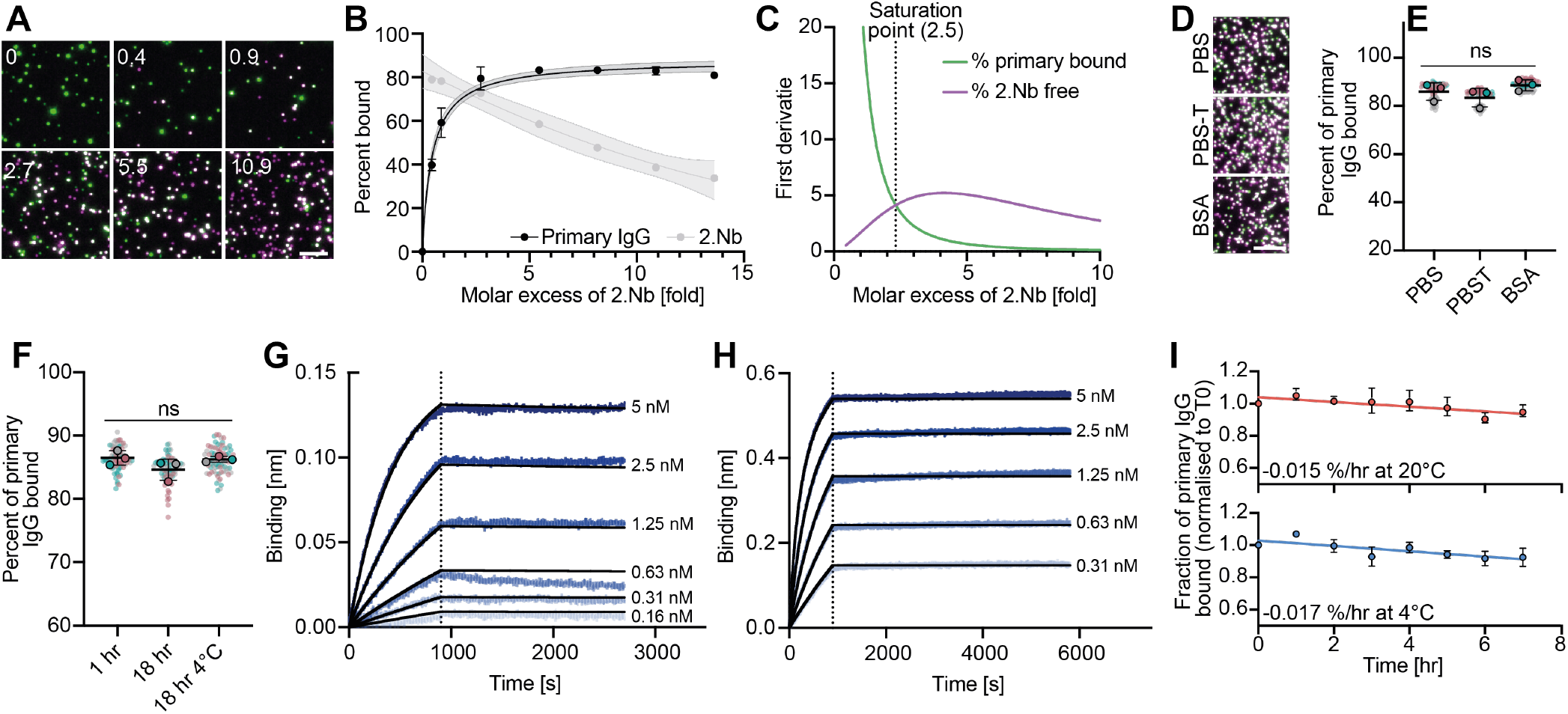
Optimized pre-incubation conditions increase fraction of primary IgG associated with 2.Nb. (**A**) TIRFM images of primary IgG (green) and 2.Nb (magenta) following 1 hr pre-incubation with the specified excess of 2.Nb. Scale bars 5 µm. Representative images shown, n = 3 with 25 technical repeats. (**B**) Scatter plot tracking the percent of primary IgG colocalized with 2.Nb in (A) fit using a Hyperbola model (black) and the percent of 2.Nb colocalized with primary IgG fit with a second-order polynomial (gray), mean ± SD of raw data shown with non-linear fit and 95% confidence intervals. (**C**) The first derivative of the fractions of bound primary IgG (green) and free 2.Nb (purple) for the data in (B). This describes the change in gradient of the fit lines in (B) as the primary IgG becomes saturated by 2.Nb. The critical saturation point was determined as the intercept of these curves, representing the point at which the increase in unwanted free 2.Nb exceeds the increase in bound primary IgG. (**D**) Representative TIRFM images of primary IgG (green) and 2.Nb (magenta) following 1 hr pre-incubation in PBS (top), PBS with 0.1% Tween 20 (middle), or PBS with 1% BSA (bottom). Scale bar 5 µm. (**E**) SuperPlot showing percent of primary IgG colocalized with 2.Nb for full dataset represented in (D). Overall statistical significance determined by one-way ANOVA. (**F**) SuperPlot showing percent of primary IgG colocalized with 2.Nb following primary IgG pre-incubation with 4.6 molar excess of 2.Nb for increasing times at room temperature (RT) or 4°C as specified. Overall statistical significance determined by one-way ANOVA, n = 3 with 25 technical repeats. (**G-H**) BLI traces of the association (left of dotted line) and dissociation (right of dotted line) of either the 2.Nb (5-0.16 nM) to immobilised primary IgG (G) or the primary IgG (5-0.31 nM) to immobilised 2.Nb (H) at 25°C. Raw data traces following baseline subtraction displayed (blue) alongside a 1:1 (E) or bivalent (F) global fit model. (**I**) Time-course data tracking the loss of bound primary IgG over time under immunoassay-relevant conditions. Time zero (T0) is the bound primary IgG fraction following 1 hour pre-incubation at room temperature, following which samples were diluted to 6.7 nM and incubated at either room temperature (top) or 4°C (bottom), sampling every hour. Mean ± SD of raw data shown with simple linear regression fit.

To explore the optimal ratio of primary IgG to 2.Nb, the fraction of primary IgG associated with 2.Nb was fit to a hyperbola and the fraction of 2.Nb associated with primary IgG fit to a second-order polynomial. This allowed the gradient of the curves at 0.01 intervals to be analyzed (the first derivatives; Figure 2C). We defined the critical saturation point as the point at which the free 2.Nb rate of increase first exceeds the bound primary IgG rate of increase, which is given by the intercept of the respective first derivative curves (Figure 2C). In the case of the 10E10 anti-rabbit IgG 2.Nb analyzed, the critical saturation point was at 2.53 molar excess of 2.Nb. At this point, 74.4 ± 1.8% of primary IgG was associated with 2.Nb. Further improvement up to 82% can be achieved with greater 2.Nb excess, but at the cost of rapidly increasing free 2.Nb that may subsequently drive non-specific interaction.

It is important to note that the accuracy of this saturation value is dependent on the accuracy with which the stock antibody concentrations are known, and this is challenging to determine absolutely. In this case it was assessed using UV-Vis spectrophotometry to measure the absorbance of the dye conjugate at its absorption maximum; the protein concentration was then obtained by correcting for the lot-specific number of dye molecules per antibody (degree of labeling, DoL) as provided by the vendor and confirmed by single-molecule photobleach analysis (Supplementary Figure 1).

To further interrogate the consistency of the derived saturation value, we used the same methodology to analyze the saturation point of most commercially available secondary nanobodies, where binding affinities spanned the low picomolar to low nanomolar range (Table 1 and Supplementary Figure 2). Interestingly, monoclonal products that have two target sites per primary antibody produced a mean saturation point at 2.43 ± 0.59 2.Nb molar excess, whereas biclonal products that have four target sites per primary antibody produced a mean at 5.04 ± 2.13. This suggests a general rule where a 20-25% 2.Nb surplus beyond the available number of binding sites is optimal and sufficient.

**Table 1.**
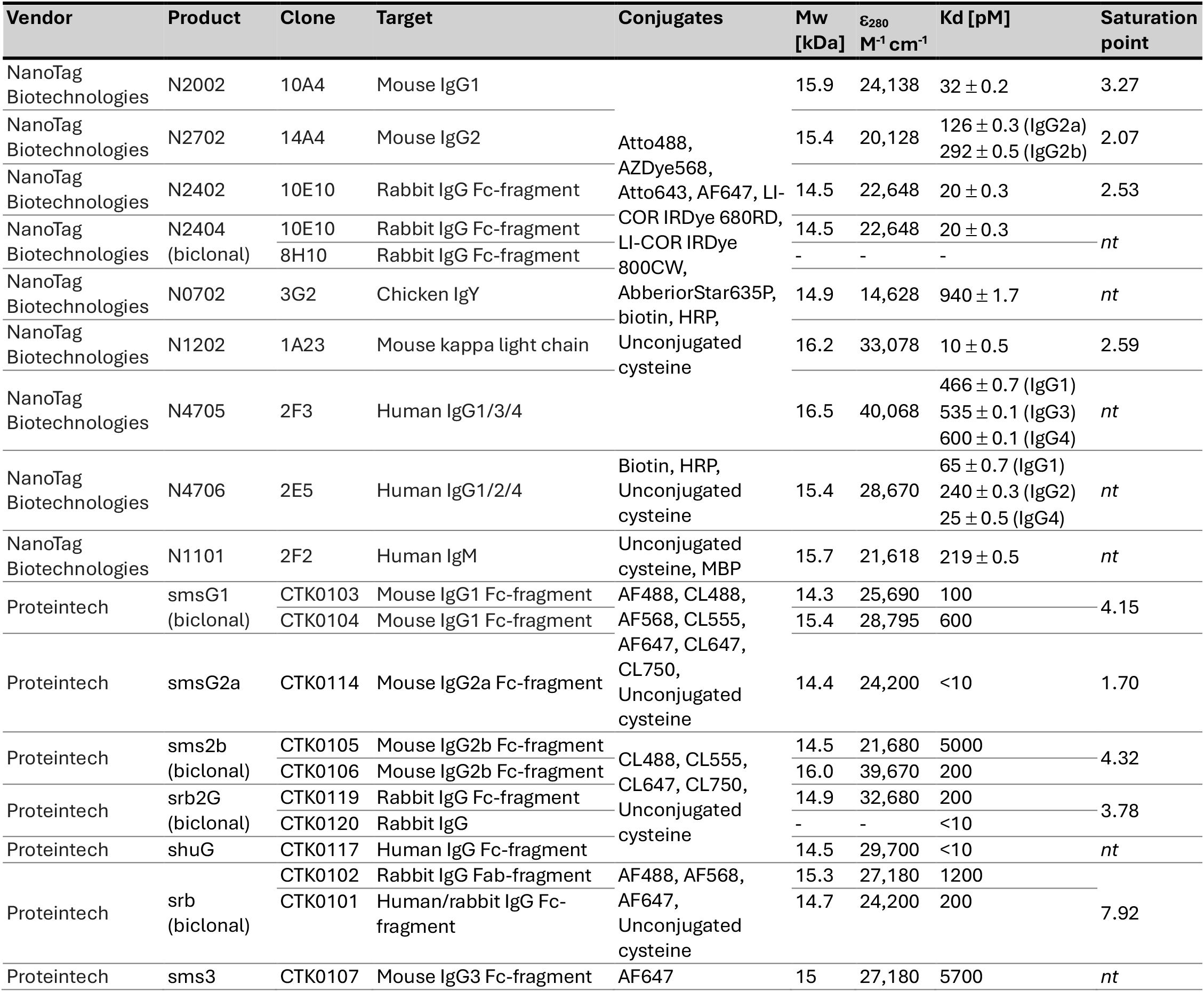
Details of commercially available secondary nanobodies and their binding properties. The 2.Nb clone, target, available conjugates, molecular weight (Mw), extinction coefficient at 280 nm (ε_280_), and Proteintech Kd data were obtained from vendor datasheets at the time of writing. More conjugates may be available as custom requests. The Kds of NanoTag Biotechnologies secondary nanobodies were determined by NanoTag Biotechnologies using Bio-layer Interferometry and provided for the compilation of this table. The saturation point (point at which the increase in bound primary IgG is exceeded by the increase in free 2.Nb) was empirically determined using our saturation assay, accompanying saturation curves are provided in Supplementary Figure 4. Not tested (*nt*) indicates cases where saturation point assessment has not been carried out.

Notably, the percentage of primary IgG associated with 2.Nb following saturation never reached 100% and varied considerably between 42-92%. We thus sought to determine if this binding efficiency could be improved by more favourable experimental conditions. 2.Nb pre-incubations must be carried out at low concentration for experiments to be economically viable, however, the effect of protein adsorption to lab plastics will be more prominent in this case. We therefore compared the binding efficiency in PBS alone to (i) PBS supplemented with 0.1% Tween 20 (PBST), a non-ionic surfactant that deters protein aggregation and surface adsorption by increasing their hydrophilicity and disrupting both hydrophobic and electrostatic interactions, and (ii) PBS supplemented with 1% bovine serum albumin (BSA), a common blocking agent that itself adsorbs to surfaces and thus blocks available binding sites (Figure 2D). No significant improvement in binding efficiency was observed over a one-hour pre-incubation using Protein LoBind tubes in the presence of either Tween 20 (83.5 ± 3.9%) or BSA (88.6 ± 2.3%) as compared to PBS (86.0 ± 3.7%) (Figure 2E). Similarly, no significant gain in binding efficiency was obtained by increasing the pre-incubation time from one hour to overnight (Figure 2F), indicative of a rapid on-rate. In addition, post-fixation of the pre-incubated mix using 2.4% paraformaldehyde, produced a plateau at 68% (Supplementary Figure 3). This suggests that the secondary nanobody has a sufficiently low off-rate as to preserve associations without fixation, with fixation perhaps inducing fluorophore quenching.

**Figure 3.**
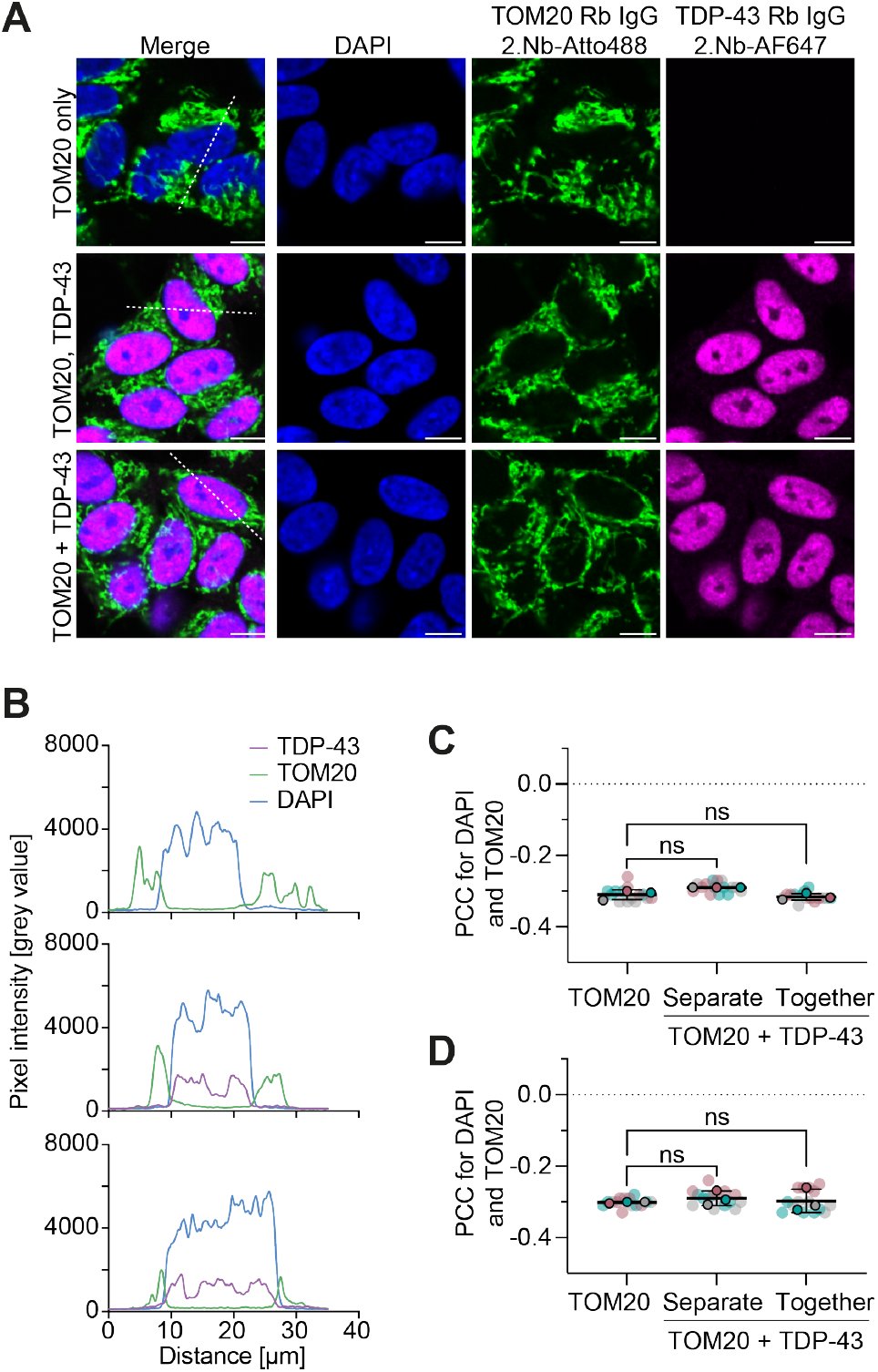
2.Nb binding is highly specific in biologically-relevant multiplexed imaging using matched species primary antibodies. (**A**) Representative spinning disk confocal images displayed as merge (left) or single channels showing fixed SH-SY5Y cells stained with rabbit anti-TOM20/ATTO488-2.Nb (green), rabbit anti-TDP-43/AF647-2.Nb (magenta), and DAPI (blue). Cells were either stained for TOM20 alone (top), TDP-43 and then TOM20 in separate staining steps (middle) or both TOM20 and TDP-43 in a single step (bottom). Scale bars 10 μm. (**B**) Line profiles of pixel intensity per channel along the white dotted line indicated in (A). (**C**) PCC analysis to assess the correlation of TOM20 with DAPI in the absence and presence of nuclear TDP-43 staining using primaries raised in the same host species at 2-fold and (**D**) 3-fold 2.Nb excess. These were assessed both when staining TDP-43 and TOM20 together and sequentially. Statistical significance assessed by Kruskal-Wallis test with post-hoc Dunn’s means comparison.

Together these findings suggest that the secondary nanobody rapidly associates, reaching equilibrium within a one-hour pre-incubation, and has a sufficiently low off-rate as to preserve associations over the course of the experiment. To verify this, binding properties of the 10E10 2.Nb clone were determined using bio-layer interferometry (BLI). The association of the 2.Nb to immobilized biotinylated primary IgG on a streptavidin-coated biosensor exhibited a k_on_ of 2.05 ± 0.004 x10^5^ M^-1^s^-1^ and a k_off_ of 9.12 ± 0.27 x10^−6^ s^-1^, producing a Kd of 20.51 ± 0.06 pM (Figure 2G). This is in strong agreement with the Kd determined by the vendor (NanoTag Biotechnologies) of 19.7 ± 0.3 pM, which is listed in comparison to all other vendor-determined binding properties in Table 1. Reversal of the ligand and analyte by immobilization of biotinylated 2.Nb, demonstrated a Kd below the 10 pM detection limit of the instrument, likely due to an avidity effect caused by two-site binding of the primary IgG (Figure 2H). Collectively, these binding kinetics support the rapid and stable association of the 2.Nb clone for primary IgG and given they lie at the lowest end of the measurable range, the true Kd may be lower than observed.

Immunoassays, however, are commonly carried out at low concentration, low temperature and for periods up to 18 hours. It is critical that the 2.Nb and primary IgG association remains stable under these conditions. To verify this, 2.Nb and primary IgG were pre-mixed and incubated one hour at room temperature, following which the “time zero” percentage of primary IgG associated with 2.Nb was determined using our single-molecule imaging approach. The pre-mix was then diluted such that the primary IgG was at the commonly used immunostaining concentration of 1 µg/mL and stored either at room temperature or 4°C, mimicking the conditions of an immunoassay. By monitoring the associated fraction at 1 hr intervals, a loss of 0.015 % per hour was observed at room temperature (Figure 2I). A paired t-test indicated that this was not significantly altered by incubation at 4°C, which produced a loss of 0.017 % per hour (Figure 2I). Extrapolating this to an 18 hour incubation produces a loss of just 0.27-0.31%, signifying that at the 20 pM affinity of the 10E10 2.Nb, binding is highly stable under experimentally-relevant conditions.

In the case that primary IgG is not fully saturated following pre-incubation or becomes available during immunostaining, there is a risk of cross-over binding of the free 2.Nb when multiplexing pre-incubations utilizing the same species of primary IgG. We therefore explored methods to deplete the free 2.Nb by size or affinity using centrifugal filtration or immuno-depletion with magnetic beads decorated with rabbit IgG. While the immuno-depletion had no apparent effect (Supplementary Figure 4A-C), spin filtration partially reduced the free 2.Nb fraction (from 32.8% ± 5.8% to 19.7% ± 3.8%), but was unable to completely clear it (Supplementary Figure 4D-F). Multiplex specificity studies are therefore critical to assess if this residual free 2.Nb drives off-target binding in immunoassays.

### Analysis of specificity for multiplexed applications

The precise concentration of commercially-sourced antibodies cannot be readily verified due to the presence of carrier protein. As “active” antibodies are unlikely to retain their starting concentration over time and with even one freeze-thaw cycle, it is critical that there is sufficient flexibility of the 2.Nb excess around the identified saturation point to accommodate these uncertainties without generating staining artefacts.

To evaluate this in a biologically-relevant assay, SH-SY5Y cells were stained with two different primary antibodies, both of which were raised in rabbit, using a 2.Nb pre-incubation. Anti-TOM20, which targets mitochondria, was pre-incubated with anti-rabbit 2.Nb labeled with ATTO488, while anti-TDP-43, which is predominantly nuclear, was pre-incubated with anti-rabbit 2.Nb labeled with AF647. If the presence of free antibody species or nanobody hopping drives mis-targeted staining, an increase in ATTO488 signal will be observed in the nucleus only when anti-TDP-43 is also present. Using this specificity assay, it was possible to quantitatively assess specificity changes when using a 2.Nb molar excess slightly above or below the 2.53 saturation point determined above.

In the case of a two-fold 2.Nb excess, ATTO488 and AF647 appear visually well segregated in confocal microscope optical sections (Figure 3A). By drawing a line profile across select cells, ATTO488 is almost completely excluded from the nuclear region delineated by DAPI signal, while AF647 is predominantly restricted to the nuclear region consistent with its small soluble cytoplasmic population (Figure 3B). To quantify this result across a large population of cells (n≥59 obtained across three different wells), Pearson’s correlation coefficient (PCC) was used (Figure 3C). A moderate anti-correlation of -0.31 ± 0.014 was observed between ATTO488 and DAPI in the absence of rabbit anti-TDP-43 IgG. Similar coefficients were observed when staining with anti-TDP-43/AF647-2.Nb prior to or alongside the anti-TOM20/ATTO488-2.Nb stain (−0.29 ± 0.0 and -0.32 ± 0.009). The Kruskal-Wallis test indicated a significant difference among the three groups, however, post-hoc Dunn’s means comparison did not reveal significant pairwise differences. This discrepancy is likely due to the lack of variability across one of the groups. Interestingly, a higher level of variation was observed when a 3-fold molar excess was used (Figure 3D), however these again presented no statistically significant differences between groups.

We finally sought to verify this by ascertaining the precise percentage of mis-targeted labeling at a single-molecule level. For this purpose, TDP-43 from SH-SY5Y cell lysate was immobilized at single-molecule density on a PEG-passivated surface using previously described protocols^19^. This was subsequently probed with rabbit anti-TDP-43 directly labeled with CoraLite 594 (CL594) and pre-incubated with anti-rabbit 2.Nb-AF647 (Figure 4A). On-target detections will thus appear as coincident CL594 and AF647 positive puncta by TIRF microscopy. In addition, a second rabbit antibody/2.Nb pair, for which no antigen was present on the surface (anti-phosphorylated alpha-synuclein, paSyn), was applied either following or alongside the anti-TDP-43 mix. In this case, the rabbit primary IgG was paired with ATTO488-labeled anti-rabbit 2.Nb. Mis-targeted 2.Nb binding events caused by multiplexing two pre-mixes of the same species can then be identified as CL594 and ATTO488 positive puncta (Figure 4A-B).

**Figure 4.**
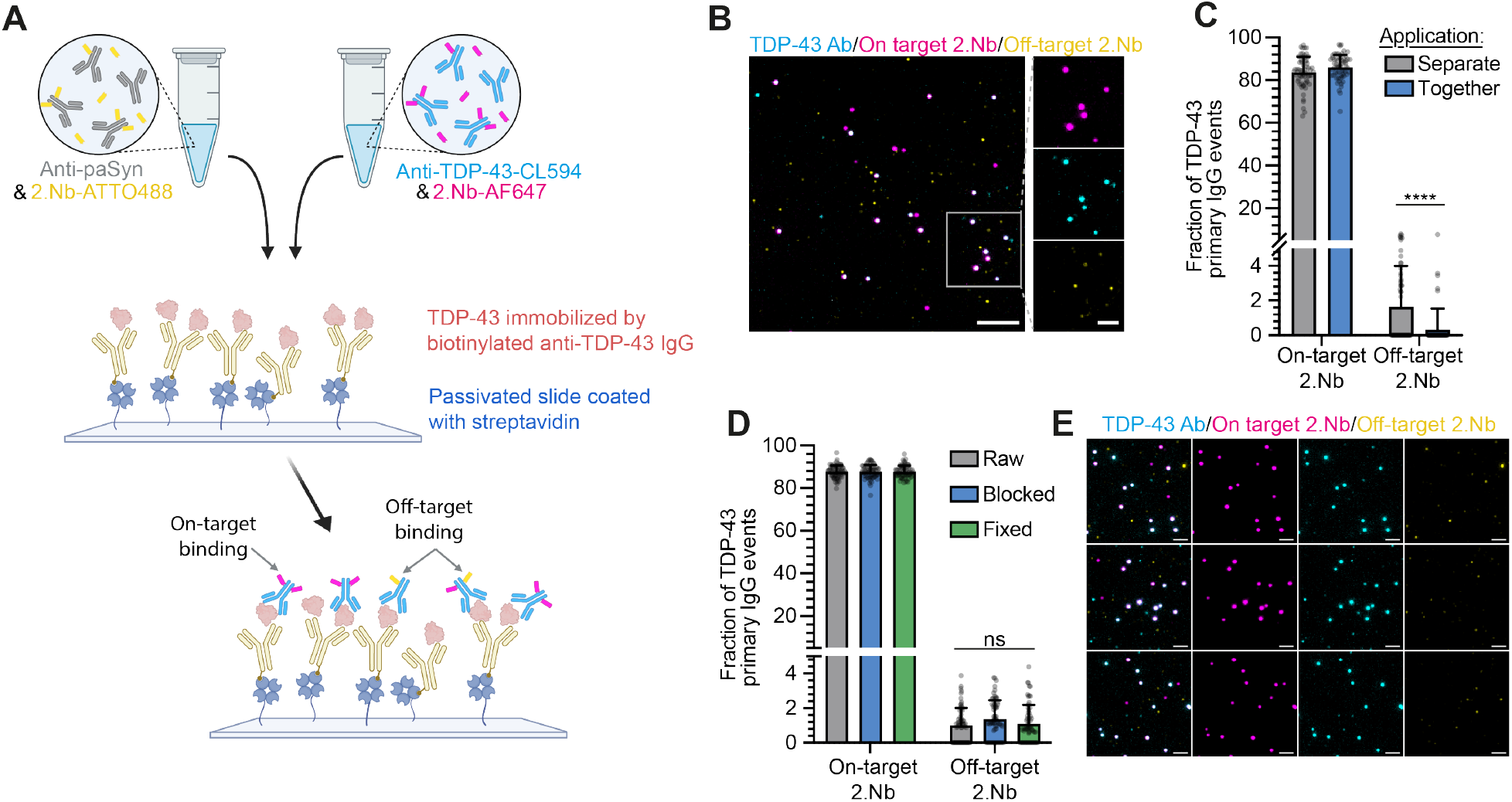
Off-target 2.Nb binding during same species multiplexing is minimal at the single-molecule level. (**A**) Schematic of the experimental design; TDP-43 (red) derived from SH-SY5Y cell lysate is immobilised on a passivated slide coated with mouse anti-TDP-43 capture antibody (yellow). The surface is then probed with rabbit anti-TDP-43 (CL594 labelled) pre-incubated with anti-rabbit 2.Nb (AF647 labelled). The availability of epitopes for cross-over binding or nanobody hopping during multiplex assays is determined by additional application of anti-rabbit 2.Nb (ATTO488 labelled) pre-incubated with rabbit anti-paSyn (for which there is no target present on the surface). (**B**) A representative TIRF microscopy image of the assay, showing rabbit anti-TDP-43 CL594 (cyan), anti-rabbit AF647 2.Nb (magenta), and anti-rabbit ATTO488 2.Nb (yellow). On-target binding is thus represented by cyan/magenta coincident events and unwanted cross-over binding by coincident cyan/yellow or cyan/magenta/yellow events. Single-channel images for the region indicated are shown (right). Scale bars 5 μm in full-field and 2 μm in image crops. (**F**) The percentage of CL594-positive spots that show on-target (AF647 coincident) and off-target (ATTO488 coincident) binding both when applying the pre-incubation mixes together and separately. Statistical analysis of off-target binding used a Mann-Whitney test (p < 0.0001). (**G**) The percentage of CL594-positive spots that show on-target and off-target binding when applied sequentially and either left untreated (gray), blocked with an excess of unlabeled 2.Nb (blue), or fixed with 4% PFA (green). Statistical significance of oL-target binding determined using a Kruskal-Wallis test. (H) Representative single channel and merged TIRF microscopy images of the data presented in (G). Scale bars 2 µm.

Using this assay, it was apparent that the percentage of mis-targeting events is extremely low, with less than 2% of CL594 detections colocalizing with off-target labels (Figure 4C). Of note, a significantly higher proportion of mis-targeted events occurred when the pre-incubated antibodies were added sequentially (1.64% ± 2.36%) rather than simultaneously (0.32% ± 1.22%). Various techniques have been proposed in the literature for the suppression of mis-targeted 2.Nb binding, including the application of an excess of unlabeled 2.Nb to saturate any remaining free binding sites on the primary antibody, or fixation to deter nanobody hopping. Using the same single-molecule assay with sequential anti-TDP-43/2.Nb-AF647 and anti-paSyn/2.Nb-ATTO488 application, these approaches were unable to significantly quench mis-targeted events (Figure 4D-E). In the absence of any intervention 1.01 ± 1.01% mis-targeted events were observed, whereas in the presence of unlabeled 2.Nb at four molar excess or 4% PFA fixation, 1.37 ± 1.08% and 1.09 ± 1.09% were observed respectively. These findings suggest that antibody hopping, which would be quenched by PFA fixation, does not contribute significantly to cross-over labeling, which is consistent with the low off rates observed above. Given the increase in off-target events upon sequentially stained samples and the lack of improvement obtained with an unlabeled excess of 2.Nb, we speculate that the wash step itself may be increasing the availability of binding competent epitopes. For experiments that demand cyclical staining, delayed or extremely long imaging times, sample fixation post-staining may be advantageous to prevent further epitope unmasking.

Together, these results suggest that 2.Nb pre-incubations carried out with 2.Nb at an excess close to its binding saturation point can produce highly specific target binding.

## Discussion

Secondary nanobodies present great potential for life sciences research, with immunoassays contributing a central methodology across research fields. Their adoption for one-step IF, where the primary IgG and secondary nanobody are pre-mixed prior to application, will simplify staining workflows, enable primary antibody selection based on binding properties rather than species type, support easy uptake of new dyes and tags, and enable highly multiplexed imaging without the cost and expertise of direct primary IgG conjugation. In addition, their pre-association with primary IgG prior to sample application may reduce non-specific staining driven by high secondary antibody concentrations and the defined degree of labeling supports improved quantification over polyclonal secondary antibodies.

However, multiplexed use of 2.Nbs with primary IgG of the same species presents an additional avenue for imaging artefacts to arise through cross-over labeling or nanobody hopping. It is therefore critical that these new tools are carefully evaluated and appropriate protocols implemented. To support this, we have described a series of straight forward imaging assays and provide accompanying analysis scripts for the optimization and validation of 2.Nb binding, spanning the single-molecule to cell level.

We used these assays to analyze the appropriate conditions for primary IgG and 2.Nb pre-incubation and have identified the required 2.Nb excess for binding saturation across commercially available 2.Nbs, which follow a general rule of 20-25% excess beyond the number of available binding sites. Overall, we found that pre-incubation in protein LoBind tubes of the primary IgG diluted in standard buffers to 10-30 nM with the appropriate excess of 2.Nb for one hour at room temperature was sufficient and optimal.

It must be noted, however, that a population of free primary IgG persists that cannot be resolved by increasing the 2.Nb concentration or altering incubation conditions. Similarly, approximately 20% 2.Nb remains non-bound when the primary IgG is instead in excess (Figure 2B). Though the cause is unknown, we posit that these may not be active or binding-competent species, perhaps due to degradation, misfolding, fragmentation, or epitope occlusion by the dye conjugate. A fraction may also represent residual free dye molecules not associated with primary IgG or primary IgG bound by unlabeled 2.Nbs caused by the fluorophore labeling processes. Indeed, the peak saturation of primary IgG associated with 2.Nb appears to be dictated by the identity of the primary antibody used. For example, the 2.Nb clones sms2b, 14A4, and 1A23, were paired with the same primary IgG and produced similar peak saturations at 87.2%, 85.3%, and 83.8% respectively, despite divergent saturation points (Supplementary Figure 2).

In agreement that these free species are not binding-competent and thus unable to drive mis-labeling, multiplex assays demonstrated <2% of primary IgG is bound by an off-target secondary nanobody. Indeed, cell-based assays demonstrated no quantitative difference in antibody localization in single versus multiplex assays.

We conclude that secondary nanobodies are a robust and powerful tool. Our validation data confirms their suitability for the quantitative evaluation of multiplexed matched species primary IgG and as a low-cost alternative to direct primary antibody labeling for use across the spectrum of immunoassays.

## Methods

### Antibodies

The product details and concentrations per experiment of all antibodies used in this study are outlined in Supplementary Table 1.

### Primary IgG and 2.Nb pre-incubation

Primary IgG and 2.Nb were combined in 0.5 mL Protein LoBind tubes (Eppendorf, 0030108094) with a minimum 5 μL volume. They were diluted as required in 0.22 μm-filtered PBS (ThermoFisher Scientific, 10010023) supplemented with 0.1% Tween 20 (PBST) unless otherwise specified. Primary IgG and 2.Nb pre-incubation concentrations were in the range of 12-500 nM and 4-1500 nM, respectively. Experiment-specific concentrations are provided in Supplementary Table 1. Pre-incubations were carried out for 60 min at room-temperature in the dark unless otherwise specified, following which they were diluted up to 2000-fold in PBST immediately prior to slide application. Experiments investigating buffer composition used carrier-free proteins and used PBS as the final diluent. Where specified, fixation was carried out by addition of 2.4% final concentration paraformaldyhde (ThermoFisher Scientific, 28906) to the pre-incubation product and incubated 7 min prior to 100x dilution and slide application.

### Single-molecule binding assay

Borosilicate glass coverslips were exposed to low-pressure argon plasma for 45 min using a Diener Electronic Zepto One Low-Pressure Plasma System to remove contaminants and activate the surface for protein adsorption. 9 mm Frame-Seal incubation chambers (Bio-Rad, SLF0201) were applied to each slide. 50 μL sample was applied and incubated for 30 min in a dark humidified chamber at room temperature, before washing with 0.22 μm-filtered PBST (or PBS for experiments investigating buffer composition). Samples were applied at staggered 2 min intervals to ensure precise incubation timings.

### Total internal reflection fluorescence microscopy

Samples were imaged in 50 μL PBS by TIRF microscopy using an ONI Nanoimager equipped with a 100x/1.4 numerical aperture oil immersion objective lens and ORCA-Flash 4.0 V3 sCMOS camera. Samples were exposed sequentially to 638 nm and 488 nm light at a 53.5° illumination angle. A 640 nm dichroic split emission onto different regions of the camera chip. TetraSpeck beads (ThermoFisher, T7279) were captured alongside each dataset to enable accurate channel alignment. Each field of view (FOV) was imaged for 10 frames at a rate of 20 frames s^-1^, and a 5 × 5 grid of 200 μm-spaced FOVs was captured per condition to account for variation.

### Single-molecule image processing and analysis

Image pre-processing and single-molecule detection was performed as previously described for single-molecule two-colour aggregate pulldowns (STAPull)^19^ using custom written ImageJ macros and python code (DOI: 10.5281/zenodo.8279273). Briefly, single-channel images were registered based on TetraSpeck ground truth data using the imreg_dft Python library^20^. Image series were maximum intensity projected and hot pixels detected and eliminated using the in-built Fiji ‘remove outliers’ function (threshold>2000, 1 pixel radius). Particle detection, colocalization, and quantification were performed using the ComDet v.0.5.5 plugin for ImageJ (https://github.com/ekatrukha/ComDet). Threshold settings were optimized to minimize false positive detections in negative control samples. Two-colour events with a centre of mass within two-pixel proximity were considered coincident. Images were visually inspected and those with uneven illumination or discernible dirt were rejected. The percentage of primary IgG labeled was computed per FOV as the fraction of coincident events to total CL488-positive events. Each biological repeat represents the mean of all analyzed FOVs.

### Derivation of 2.Nb binding properties

The fraction of (i) primary IgG bound to 2.Nb and (ii) 2.Nb bound to primary IgG were plotted as a function of 2.Nb molar excess using GraphPad Prism v10.4.1. Standard curves were fit using a hyperbola model for the fraction of bound primary IgG and a second-order polynomial for the fraction of bound 2.Nb. The first derivative was computed at 0.01 increments of 2.Nb excess. The ‘saturation point’ was identified from the first derivatives as the 2.Nb excess at which the magnitude of loss in bound 2.Nb first exceeds the gain in bound primary IgG. The ‘maximum bound fraction’ represents the mean fraction of bound primary IgG empirically observed above the saturation point.

### Single-molecule photobleach assay

Each antibody was prepared at 20-40 pM in 0.22 μm-filtered PBS supplemented with 0.1% Tween 20 and adsorbed to cover glass as described for the single-molecule binding assay. The intensity of emitted fluorescence was tracked across nine FOVs for 7.5 s and 30 s for AF647-labeled 2.Nb and CL488-labeled primary IgG, respectively, using TIRF microscopy as described above.

### Single-molecule photobleach analysis

Photobleach analysis was adapted from published methods^21^ using automated python scripts (DOI: 10.5281/zenodo.14945283). Single molecules were segmented from maximum intensity projections pre-processed to remove outliers, minimize noise with a gaussian filter and suppress background using a tophat filter. Single molecules were subsequently classified as features 4.5 standard deviations above the image mean. Single molecule traces were extracted from the raw image series and Chung-Kennedy filtering performed to reduce trace noise while preserving data discontinuities. Traces were filtered to reject likely protein clusters, high noise traces, and incompletely bleached events. Step change events were then detected using the pelt algorithm in the Ruptures library^22^.

### Determination of antibody concentration

The conjugate dye concentration was measured by spectrophotometric analysis using a ThermoFisher Scientific NanoDrop One. To obtain the antibody concentration, this value was subsequently corrected for the lot-specific degree of labeling specified by the vendor. The concentrations of AF647 and CL647 Plus were determined by absorbance at 650 nm and 654 nm, respectively, using extinction coefficients of 239,000 M^-1^cm^-1^ and 250,000 M^-1^cm^-1^. The concentration of AF488 and CL488 Plus were determined by absorbance at 494 nm and 493 nm, respectively, using extinction coefficients of 71,000 M^-1^cm^-1^ and 80,000 M^-1^cm^-1^.

### Bio-layer interferometry

BLI assays were conducted using a Sartorius Octet R8 system at 25°C with 1,000 rpm agitation. Streptavidin SAX biosensors (Sartorius, 18-5117) were hydrated in 1x Kinetics Buffer (Sartorius, 18-1105) for a minimum of 10 mins prior to use. Details of antibody and 2.Nb pairs and experimental timings are given in Supplementary Table 3. Kinetic parameters were derived from global fitting of binding curves using a 1:1 or 2:1 model dependent on the number of binding sites on the analyte.

### 2.Nb spin filtration

30 kDa MWCO centrifugal filters (Merck, UFC503024) were pre-rinsed with 0.22 μm-filtered PBS supplemented with 0.1% Tween 20 for 5 min at 14,000 xg at 4°C. 40 μL of primary IgG (20 nM), 2.Nb (30 nM), or a 1 hour pre-incubated mix of both (20 nM and 66 nM respectively) was loaded, diluted to 500 μL with PBST and spun at 14,000 xg for 5 min. The dilution and spin were repeated once further. The membrane was rinsed with 40 μL PBST and the sample eluted by inversion of the column and a gentle 1,000 xg 2 min spin. The flow-through and filtrate were collected in pre-weighed tubes and the volumes adjusted to match by weight such that the total sample dilution broadly matched per fraction.

### Immuno-depletion

Up to 100 μg magnetic beads coated with rabbit (DA1E) IgG (Cell Signaling Technology, 8726) were added to 20 μL primary IgG (26.5 nM), 2.Nb (90 nM), or a 1 hour pre-incubated mix of both. These were incubated for 30 min at room temperature with agitation. Beads were then separated from the sample using a DynaMag-2 Magnet (ThermoFisher Scientific, 12321D).

### Cell culture

SH-SY5Y cells were cultured in DMEM/F-12 with GlutaMAX (ThermoFisher Scientific, 31331028) supplemented with 10 % foetal bovine serum (ThermoFisher Scientific, A5670402) and 1% penicillin-streptomycin (ThermoFisher Scientific, 15070063). Cells were incubated at 37°C with 5% CO_2_ and passaged every 2-3 days using 0.5 mM EDTA.

### Immunostaining of cells

SH-SY5Y cells were seeded on a No. 1.5 polymer coverslip-based 96-well plate with IbiTreat coating (Ibidi, 89626) at a density of 30,000 cells per well and cultured overnight. Cells were rinsed with PBS three times then fixed with 4% paraformaldehyde in PBS for an initial 10 min at 4°C followed by 20 min at room temperature. Cells were washed with PBS then permeabilized by 5 min incubation in TBS (50 mM Tris pH 7.4, 150 mM NaCl) supplemented with 0.5% Triton X100 and washed with TBS. Cells were subsequently treated 1 hour with blocking buffer (3% BSA, 0.3 M glycine in TBS) prior to application of pre-incubated antibody/nanobody mix, prepared as above at either 1:2 or 1:3 primary IgG to 2.Nb. Wells were stained in triplicate with either (i) rabbit anti-TOM20 (Abcam, ab186735) pre-incubated with ATTO488 2.Nb (NanoTag Biotechnologies, N2402-At488-S), (ii) rabbit anti-TDP-43 (Proteintech, 12892-1-AP) pre-incubated with AF647 2.Nb (Nanotag Biotechnologies, N2402-AF647-S), or (iii) a pooled mix of both. Details of pre-incubation and final concentrations are provided in Supplementary Table 1. Cells were covered and incubated 1 hour at room temperature, then washed three times with TBST (TBS with 0.1% Tween 20). A freshly prepared pre-incubation of anti-TOM20 and ATTO488 2.Nb was then applied to all wells stained with anti-TDP-43, incubated 1 hour at room temperature then washed three times with TBST. All wells were washed a further three times with TBS then stained 10 min with 2 μg/mL DAPI. Wells were washed three times with TBS, then imaged immediately.

### Cell imaging

Cells were imaged on a Revvity Opera Phenix Plus High-Content spinning disk onfocal microscope equipped with a 40x/1.1 NA water objective lens, producing an optical section thickness of 1.2 µm. Image stacks were captured at 1 µm Z-interval with 0.149 μm pixel size using two Andor Zyla OEM sCMOS cameras. To detect DAPI fluorescence, the sample was excited at 375 nm for 100ms and emission between 435-480 nm collected. To detect ATTO488, the sample was excited at 488 nm for 300 ms and emission between 500-550 nm detected. To detect AF647, the sample was excited at 640 nm for 400 ms and emission between 650-760 nm collected. Dual camera images were auto-aligned post-acquisition using the Harmony software package (Revvity).

### Pearson’s correlation coefficient analysis

PCC analysis was carried out using a custom-written ImageJ macro (DOI: 10.5281/zenodo. 14945331). The most focussed slice of each image stack was identified and extracted based on DAPI signal intensity. Both segmentation and colocalization analysis were carried out in this slice. Total cell area was segmented from a summed image of DAPI and TOM20 signal processed with a gaussian blur with 4 pixel sigma, a rolling ball background subtraction with 250 pixel kernel and set to high contrast. Segmentation utilized the Li autothreshold. This ROI was used to exclude background pixels from the PCC analysis carried out using the Coloc 2 plug-in for Fiji.

### Cell lysate preparation

Cell lysate preparation was based on published protocols for the isolation of non-pathological TDP-43^23^. Briefly, HS buffer was freshly prepared: 10 mM Tris pH 7.5, 150 mM NaCl, 0.1 mM EDTA, 1 mM DTT supplemented with cOmplete protease inhibitor cocktail (Merck, 11836170001) and PhosSTOP (Merck, 4906845001) tablets. SH-SY5Y cells were grown to 80% confluence on a 6-well culture plate. Cells were rinsed with PBS, then incubated 5 min with 100 μL/well lysis buffer at room temperature (2 mM MgCl_2_, 1 unit/μL benzonase, and 0.5% sarkosyl in HS buffer). All cells were harvested and transferred to a single protein LoBind tube using a cell scraper. The plate was rinsed with a further 100 μL lysis buffer, which was added to the lysate. The lysate was vortexed briefly and 4.25% sarkosyl in HS buffer added as needed to bring final sarkosyl concentration to 2%. Lysates were incubated on ice for 45 min, vortexing every 10 min, then centrifuged at 21,200 xg at 4°C. The supernatant was recovered, aliquoted and flash frozen for storage at -80°C.

### SiMPull assay

A single-molecule pull-down surface was prepared as previously described^19^. Briefly, glass coverslips (CellPath, SAN-2460-03A) were treated with argon plasma for 45 min to remove contaminants, then submerged in 0.22-μm–filtered 1 M KOH for 20 min to activate the surface for silane functionalization, rinsed in methanol, and transferred to 1% (3-aminopropyl) trimethoxysilane (Sigma-Aldrich, 281778) in methanol supplemented with 5% acetic acid for a 20 min covered incubation. The coverslips were serially washed in methanol and deionized water, blast dried with argon gas and affixed to an 18-well gasket (Ibidi, 81818). Freshly prepared 100 mg/mL PEG solution (mPEG-Succinimidyl Varelate (SVA) 5000 Da, Laysan Bio) containing 5% biotin-mPEG-SVA (5000 Da, Laysan Bio), and 0.1 M NaHCO_3_ was applied and incubated 6 hours in the dark at room temperature. The surface was then rinsed with deionized water and blast dried with argon gas, before incubating 10 min with 0.2 mg/mL streptavidin (ThermoFisher Scientific, 21125) in 0.02-μm–filtered T50 buffer (10 mM Tris pH 8.0 supplemented with 50 mM NaCl) and subsequently washed three times in T50. 100 nM biotinylated mouse anti-TDP-43 (Novus, NBP1-92695B) was incubated on the surface for 20 min, following which the surface was washed with PBS and exposed to neat cell lysate overnight at room temperature for TDP-43 capture and immobilization. Following a PBS wash, specific and non-specific rabbit primary IgG/2.Nb pre-incubations were applied (see Supplementary Table 1 for antibody details), either in combination or successively, for 20 min at room temperature. Slides were washed with PBS and imaged immediately by TIRF microscopy as described above.

Where fixation is specified, 4% PFA in PBS was applied to wells for 10 min after each antibody application had been removed and washed. Where blocking with unlabeled 2.Nb is specified, unlabeled anti-rabbit 2.Nb (NanoTag Biotechnologies, N2405-250ug) was added to the pre-incubation mix following 1 hour incubation at a final concentration of 120 nM (equivalent to 4 molar excess). This was incubated 5 min at room temperature before diluting as appropriate and applying to the surface.

### SiMPull image processing and analysis

The single-molecule image processing and analysis described above was used, adapting as required for three colour coincidence detection of Atto488, CL594, and AF647 puncta (DOI: 10.5281/zenodo.14945338). The ComDet detection in this case required single molecule detections to satisfy a minimum particle size of 4 pixels and have an intensity 20 SD above the image mean. For each FOV, the CL594 coincidence with ATTO488, AF647, and both ATTO488/AF647 was determined for the test sample and a control image where the AF647 channel was flipped vertically and the CL594 channel flipped horizontally with respect to the others to create an offset between all channels for computation of ‘chance’ coincidence. The total CL594/AF647 events (Σ_*specific*_) were used to compute the percentage of anti-TDP-43 with on-target 2.Nb bound as follows:

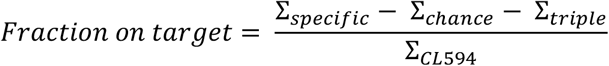

Where Σ_*chance*_ is the chance event contribution, Σ_*triple*_ the triple coincident events, and Σ_*CL*594_ the total CL594 positive events. Similarly, the fraction of off-target 2.Nb binding events was determined by:

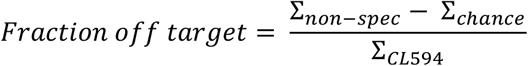

Where Σ_*non-spec*_ is the number of CL594/Atto488 coincident events (include both double and triple coincidence).

### Statistical analysis

Graphs were produced in GraphPad Prism v10.4.1. To fully convey the distribution of the data, SuperPlots were created as published^24^, presenting the mean of each biological repeat (bold) superimposed on a beeswarm plot of all technical repeat datapoints (n=25; semi-translucent) with paired colour coding. The mean and standard deviation (presented on the plot) as well as all statistical analyses were determined from the biological repeat data (n=3). Statistical significance testing was carried out in R Studio v2022.02.3 or GraphPad Prism. A Shapiro-Wilk test and Bartlett test were used to confirm normal distribution and equal variance, respectively. Tests for statistical significance were selected based on this.

### Software

Illustrations were prepared using Adobe Illustrator version 29.3.1 or BioRender (Figure 4A).

## Acknowledgements

The single-molecule instruments used in this study were funded by the UK Dementia Research Institute, UCB Biopharmaceuticals, and a donation from Dr Jim Love. The Opera Phenix Plus used in this study is supported by the Institute for Regeneration and Repair High-Content Screening Facility at the University of Edinburgh and technical assistance kindly provided by Dr Justyna Cholewa-Waclaw.

## Author contributions

Conceptualization: R.S.S. and M.H.H. Methodology: R.S.S. Investigation: R.S.S., J.O., R.Fet al., C.T.A. Visualization: R.S.S. Supervision: M.H.H. Writing – original draft: R.S.S.

## Data and materials availability

All data and software needed to evaluate the conclusions in the paper are present in the Supplementary Materials and/or shared via data repository. Raw data (DOI: 10.5281/zenodo. 14945366), single-molecule detection code (DOI: 10.5281/zenodo.8279273), Pearson’s correlation coefficient analysis code (DOI: 10.5281/zenodo. 14945331), single-molecule photobleaching analysis code (DOI: 10.5281/zenodo.14945283), three-channel coincidence analysis code (DOI: 10.5281/zenodo. 14945338).

