## Supplementary Figures for "Single-molecule validation and optimized protocols for the use of secondary nanobodies in multiplexed immunoassays"

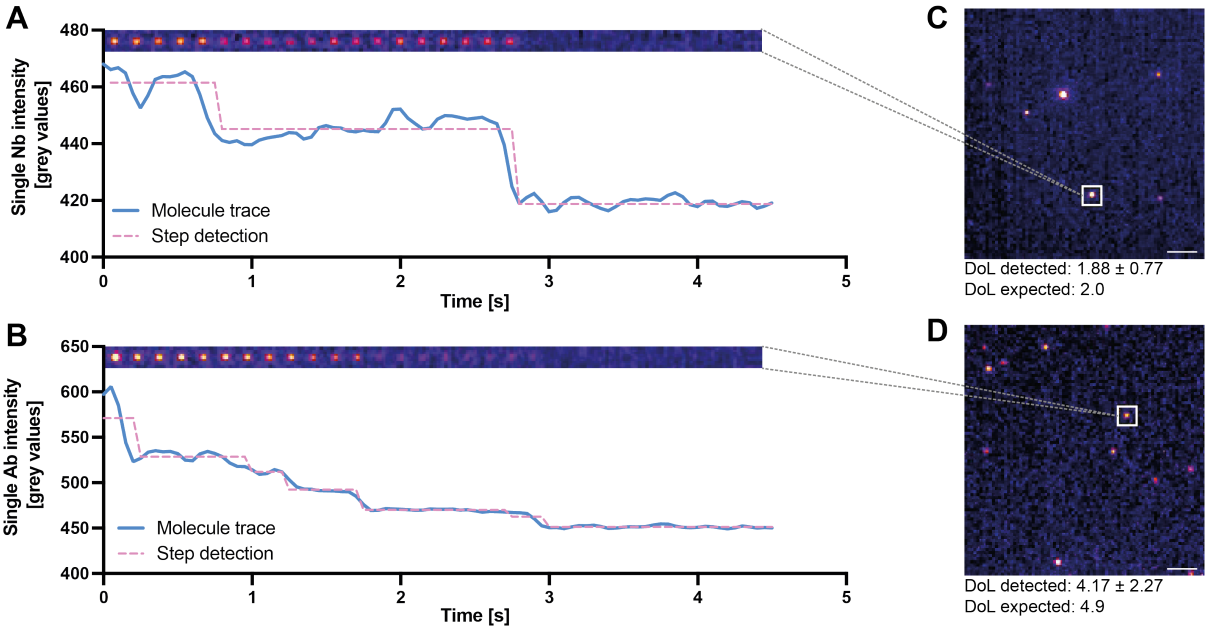


**Supplementary figure 1 – Confirmation of antibody degree of labelling using single-molecule photobleaching.** (**A-B**) Representative intensity traces for a single 2.Nb (A) or single primary IgG (B) molecule following Chung-Kennedy filtering to reduce noise (blue line) and detected step changes representative of single fluorophore bleach events (pink dotted line). Images were captured at 20 Hz and associated Gaussian-filtered raw images from 150 ms intervals are shown above each trace. (**C-D**) Representative 90-frame maximum intensity projections of TIRF microscopy images. The molecules presented in (A) and (B) are highlighted in each case. Scale bars are 2 μm. Expected and calculated degree of labelling (DoL) indicated below each image. DoL calculation based on 730 and 294 detected molecules for 2.Nb and primary IgG, respectively.

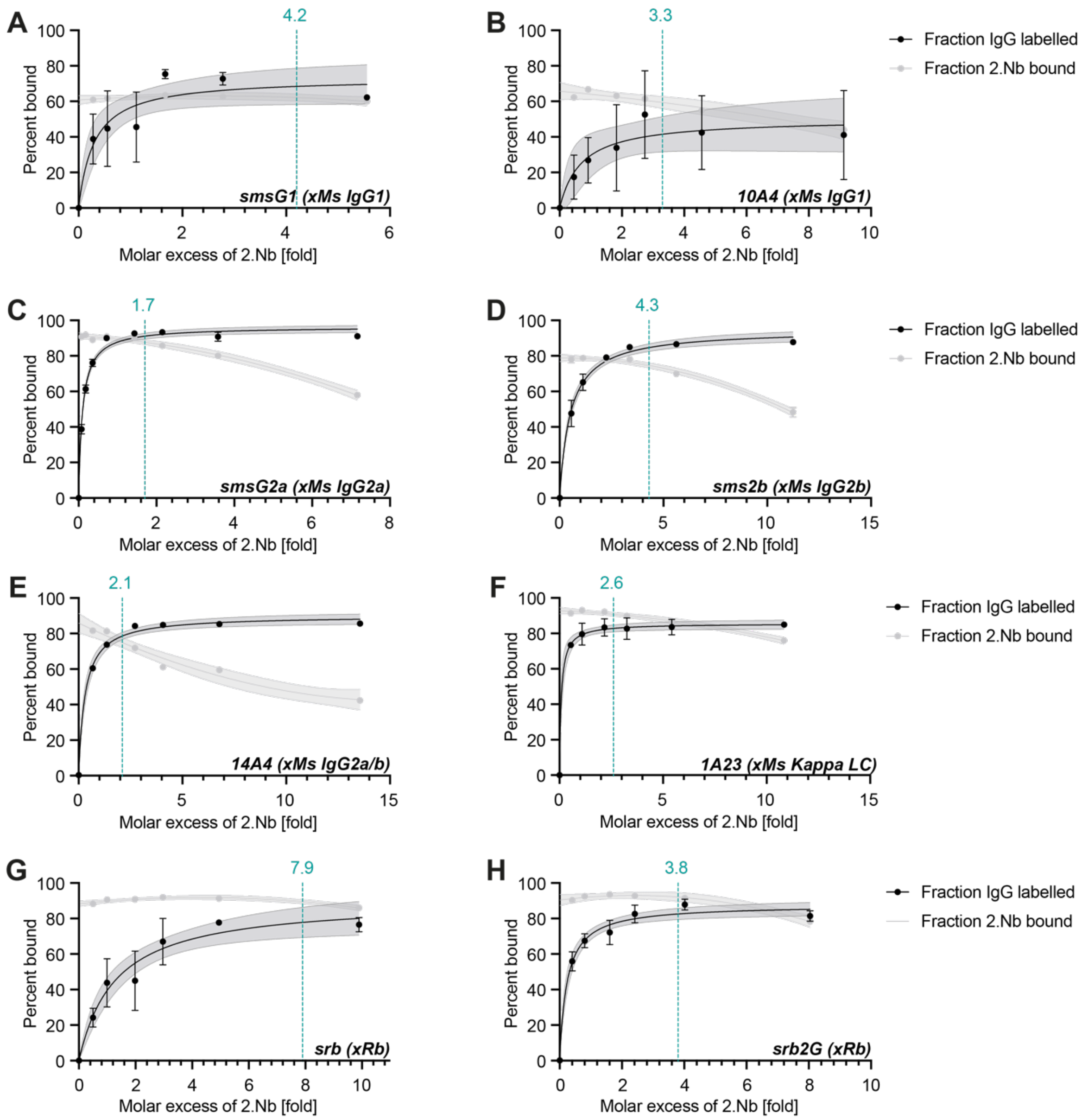


**Supplementary figure 2 – Determination of the saturation point for a panel of commercially-available 2.Nbs.** The AF647 or CL647-labelled 2.Nb indicated on each graph and further detailed in Table 1 was pre-incubated for 1 hr at room temperature with an appropriate CL488 or AF488-labelled primary IgG as described in Supplementary Table 1. The fraction of bound primary IgG (black) and bound 2.Nb (gray) was determined using TIRF microscopy following adsorption to argon plasma-treated glass. The bound primary IgG data was fit using a Hyperbola model and the bound 2.Nb data fit with a second order polynomial in all cases. The critical saturation point indicated on each graph was determined as the point at which the first derivative of the free 2.Nb exceeds that of the bound primary IgG, indicating that the fraction of free 2.Nb is growing faster than that of the bound primary IgG.

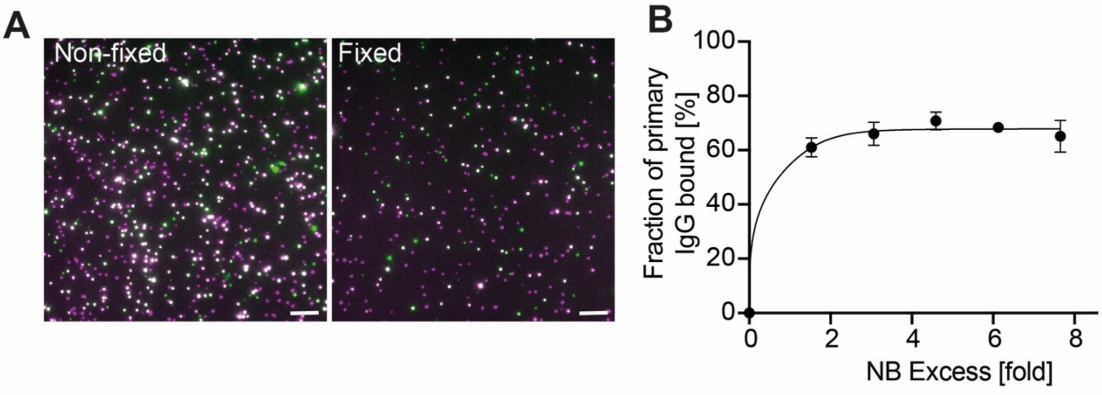


**Supplementary figure 3 – Post-fixation does not increase fraction of primary IgG associated with 2.Nb.** (**A**) Representative TIRF microscopy images of primary IgG (green) and 2.Nb (magenta) following 1 hour pre-incubation using a 4.6 molar excess of 2.Nb, shown with (right) or without (left) subsequent 7 min fixation with 2.4% paraformaldehyde. Scale bars are 5 μm. (**B**) Scatter plot tracking the percent of primary IgG colocal with 2.Nb following fixation for dataset represented in (A), mean ± SD shown with non-linear fit.

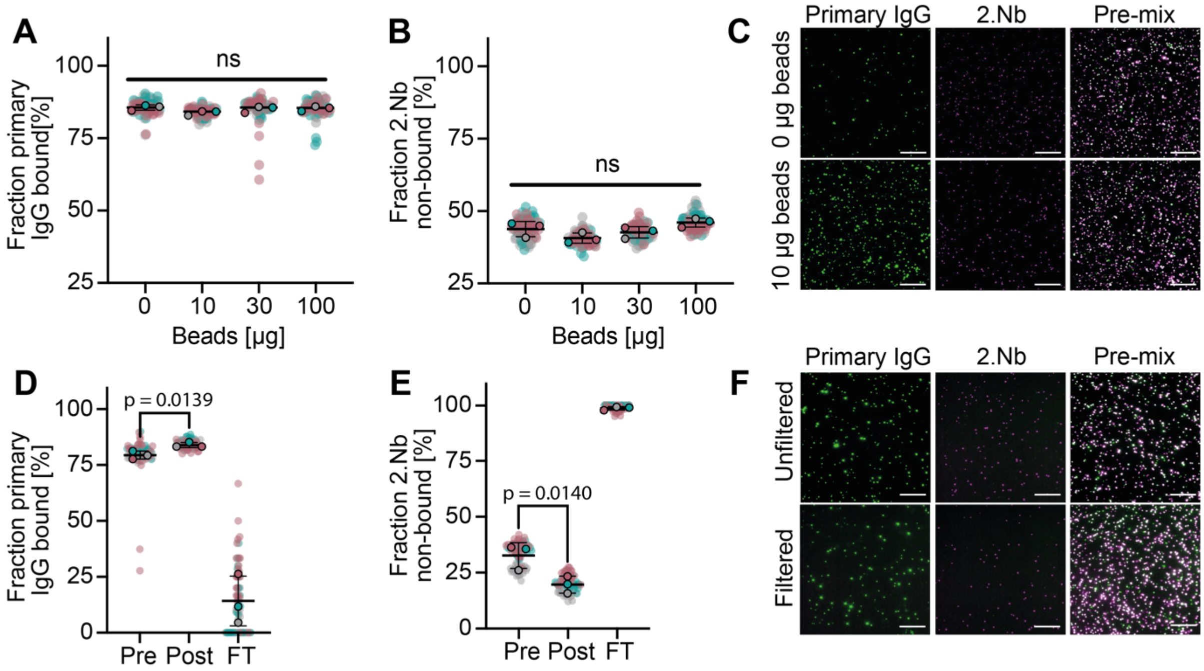


**Supplementary figure 4 – Centrifugal filtration but not immuno-depletion can reduce free 2.Nb fraction following pre-incubation.** (**A-B**) Quantification of the fraction of (A) primary IgG CL488 colocal with 2.Nb AF647 and (B) the fraction of 2.Nb AF647 only events following 1 hour pre-incubation using a 2.Nb molar excess of 3.4 and subsequent 30 min immuno-depletion of non-bound 2.Nb using increasing quantities of magnetic beads coated with rabbit IgG (as indicated). Statistical significance determined using a one-way ANOVA. (**C**) Representative TIRF microscopy images of primary IgG alone, 2.Nb alone, or the pre-incubated mix quantified in (A-B) when exposed to 0 µg or 10 µg beads as indicated. Scale bars are 10 µm. (**D-E**) Quantification of the fraction of (D) primary IgG CL488 colocal with 2.Nb AF647 and (E) the fraction of 2.Nb AF647 only events following 1 hour pre-incubation using a 2.Nb molar excess of 3.4 and either prior to (pre) or following (post) filtration through a 30 kDa centrifugal filter. Statistical significance was determined using a paired t-test to compare composition pre and post filtration. Composition of the flow-through (FT) is also shown. (**F**) Representative TIRF microscopy images of primary IgG alone, 2.Nb alone, or the pre-incubated mix quantified in (D-E) before and after filtration. Scale bars are 10 µm.
